# MissAlignment Teaches Itself Better Cryo-ET Tilt-Series Alignment by Making It Worse

**DOI:** 10.64898/2026.04.29.721716

**Authors:** Marten L. Chaillet, Joyce van Loenhout, Miguel R. Leung, Alister Burt, Dimitry Tegunov

## Abstract

Cryo-electron tomography (cryo-ET) visualizes cellular environments in a near-native state at macromolecular resolution. Accurate alignment of the tilted projection images is essential for data interpretation, yet existing reference-free algorithms often fail due to limited information overlap between images and inaccurate assumptions about the sample. Human experts, however, can easily recognize the misalignment left by these tools. We introduce MissAlignment, a machine learning approach that trains similar intuition to improve tilt-series alignment. A convolutional neural network learns to score alignment accuracy using a contrastive loss that does not require well-aligned ground truth, and gradient back-propagation from this score optimizes individual image alignment parameters. MissAlignment significantly outperforms reference-free techniques, rivals reference-based alignment, and improves all downstream analyses, making cryo-ET applicable to a broader range of biological samples.

## Introduction

Cryo-electron tomography (cryo-ET) acquires series of projection images of a biological sample at different tilt angles in a transmission electron microscope. These images are computationally aligned and backprojected into a three-dimensional reconstruction (*tomogram*). Tomograms reveal the native spatial arrangement of organelles and macromolecular complexes. When multiple copies of a complex are present, subtomogram averaging can yield high-resolution structures.

Reconstruction quality depends critically on accurate tilt-series alignment. With calibrated stage tilt and axis orientation, the sample can be treated as a rigid body whose only free parameters are 2D shifts correcting stage tracking errors. However, biological samples undergo beam-induced local deformations ^1^, violating the rigid-body assumption. While rigid-body models have achieved resolutions as high as 3.4 Å in favorable cases ^2;3^, reaching high resolution generally requires modeling non-rigid sample deformation ^4–7^.

Tilt-series alignment algorithms fall into three categories: fiducial-based, fiducial-free, and reference-based. Fiducial-based methods, dating to the earliest electron tomography work ^8^, track high-contrast markers such as colloidal gold beads. These assume fiducials are rotationally invariant and faithfully track both global and local sample motion ^9^—neither assumption holds at high resolution. Combined with the difficulty of distributing fiducials uniformly—or at all, as in cellular lamellae—this approach is increasingly impractical for modern cryo-ET.

Fiducial-free methods rely on intrinsic sample features. The simplest variant “cosine-stretches” tilt images to match adjacent projections before cross-correlation. This works well for very thin samples but degrades with increasing thickness. Because it is fast and tolerant of large shifts, it is commonly used to generate coarse initial alignments for more accurate methods such as patch tracking and projection matching.

*Patch tracking* treats the sample as a grid of virtual fiducials ^10–12^. Patches extracted from the 0°-tilt image are matched to adjacent tilts by cross-correlation and tracked pairwise to build trajectories, which are fitted to a model of fiducial Z-positions and image alignments. *Projection matching*, introduced in Protomo ^13^ and popularized by AreTomo ^14;15^, iteratively refines alignment by correlating experimental images with re-projections of an intermediate reconstruction, thereby incorporating information from non-adjacent tilts and naturally enforcing correct anisotropic filtering. A recent variant jointly optimizes alignment parameters and an implicit neural representation of the sample, though quantitative validation on experimental data is lacking ^16^.

A fundamental limitation of fiducial-free methods is the sparse information overlap between tilt images, reflected in large gaps between Fourier-space projection slices. Even where information does overlap, the relationship between tilt images is non-trivial. These methods therefore perform well on ideal samples but poorly on others. Projection matching, for instance, progressively aligns from the 0° reference to higher tilts, biasing it toward features uniform along Z (Fig. S1a). Patch tracking requires spherical particles spaced sparsely enough to remain separated at high tilt, and fails when objects overlap (Fig. S1b).

Reference-based methods, the focus of recent development, start from an initial tilt-series alignment and annotated particles to generate a low-resolution 3D reference. Alignment and particle poses are then iteratively refined by maximizing cross-correlation between reference projections and image data under geometric constraints ^4–7;17^. The known particle shape provides a strong prior linking information across tilts; as alignment improves, higher-resolution references enable progressively finer refinement. With sufficient particles, highly accurate local alignments can be obtained.

Despite offering the highest accuracy, reference-based methods face significant practical limitations. Poor initial alignment restricts convergence to large, distinctive particles such as ribosomes. When the target of interest is small, alignment can be bootstrapped from larger particles, but analysis of the small target may still suffer from local deformations if the two populations are not co-located. Moreover, the prerequisite processing adds substantial time and effort before analysis can begin.

Here, we introduce MissAlignment, a fundamentally new approach to tilt-series alignment. Rather than relying on fiducial markers, sample-specific features, or macromolecular references, MissAlignment optimizes alignment parameters by minimizing a misalignment score produced by a small 3D convolutional neural network (CNN) trained entirely without supervision. Through contrastive training using only synthetic misalignments, the network learns general indicators of alignment quality. This feature-agnostic design enables MissAlignment to generalize across diverse biological samples, approaching the accuracy of reference-based refinement while requiring no preliminary particle analysis, no manual intervention, and a fraction of the processing effort.

## Results

### Overall design

Existing alignment algorithms optimize well-defined objective functions, yet human experts can often identify residual misalignment in tomograms after these algorithms have converged. This suggests that indicators of alignment quality exist beyond what classical metrics capture. We therefore train a 3D CNN to predict a scalar misalignment score directly from tomographic reconstructions (Fig. 1a).

**Figure 1:**
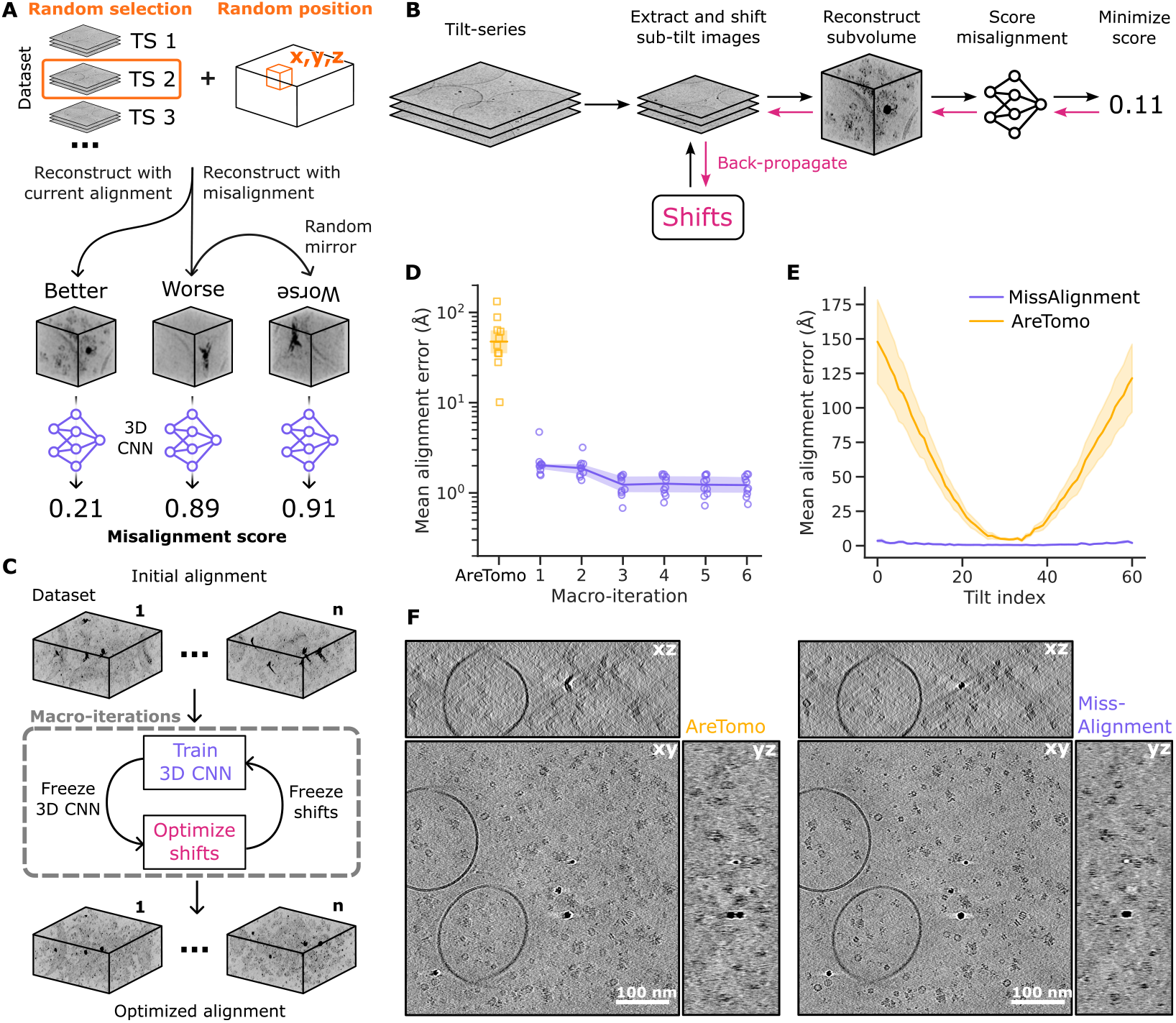
MissAlignment improves tilt-series alignment over several macro-iterations by minimizing a self-supervised contrastive misalignment score. (A) During training, a random tilt-series and 3D position are selected for reconstruction. A “better” example uses the current alignment; a “worse” example adds perturbations. Random mirroring finalizes the triplet. The loss trains the model to rank well-aligned data below poorly aligned data. (B) After training, the misalignment score is minimized by gradient descent on the image shifts through a differentiable reconstruction pipeline. (C) MissAlignment alternates training (A) and alignment over several macro-iterations. (D) Mean alignment error versus ground truth in SHREC’2119. Points show individual tilt-series; shaded area, 95% CI of the mean over all 10 tilt-series. (E) Mean alignment error as a function of tilt angle (−60° to +60°, 2° increments) for the final macro-iteration. Shaded area, 95% CI. (F) Central xy-, xz-, and yz-slices of tomographic reconstructions after AreTomo and Miss Alignment.

Because no ground-truth alignment exists, training is unsupervised and starts from misaligned data. We use a contrastive triplet loss ^18^: each triplet consists of “better” and “worse” alignment examples, produced by perturbing the image shift parameters of the current alignment (Fig. S2). This encourages the network to learn a general embedding of alignment quality.

After training, the model weights are frozen and a fully differentiable reconstruction pipeline provides gradients for the tilt image shifts with respect to the misalignment score (Fig. 1b), which are optimized by gradient descent. The scoring function can extrapolate beyond the initial alignment quality, but multiple bootstrapping iterations of training and alignment are needed to reach the best result (Fig. 1c,d). On simulated data without local deformation, MissAlignment reduces mean alignment error by an order of magnitude compared to AreTomo (Fig. 1e,f).

Training is performed from scratch on each dataset because *in situ* data can have very different statistical properties that may compromise a model pretrained on limited examples. This unsupervised, robust procedure makes MissAlignment suitable for integration into automated cryo-ET processing pipelines.

### Projection model for local alignment

MissAlignment corrects beam-induced motion through an extensible projection model that maps each 3D sample position to its corresponding 2D position in each tilt image, defining forward and backward projection operations (Fig. 2a). Additional terms describing spatially and temporally resolved deformation can be incorporated to model beam-induced motion.

**Figure 2:**
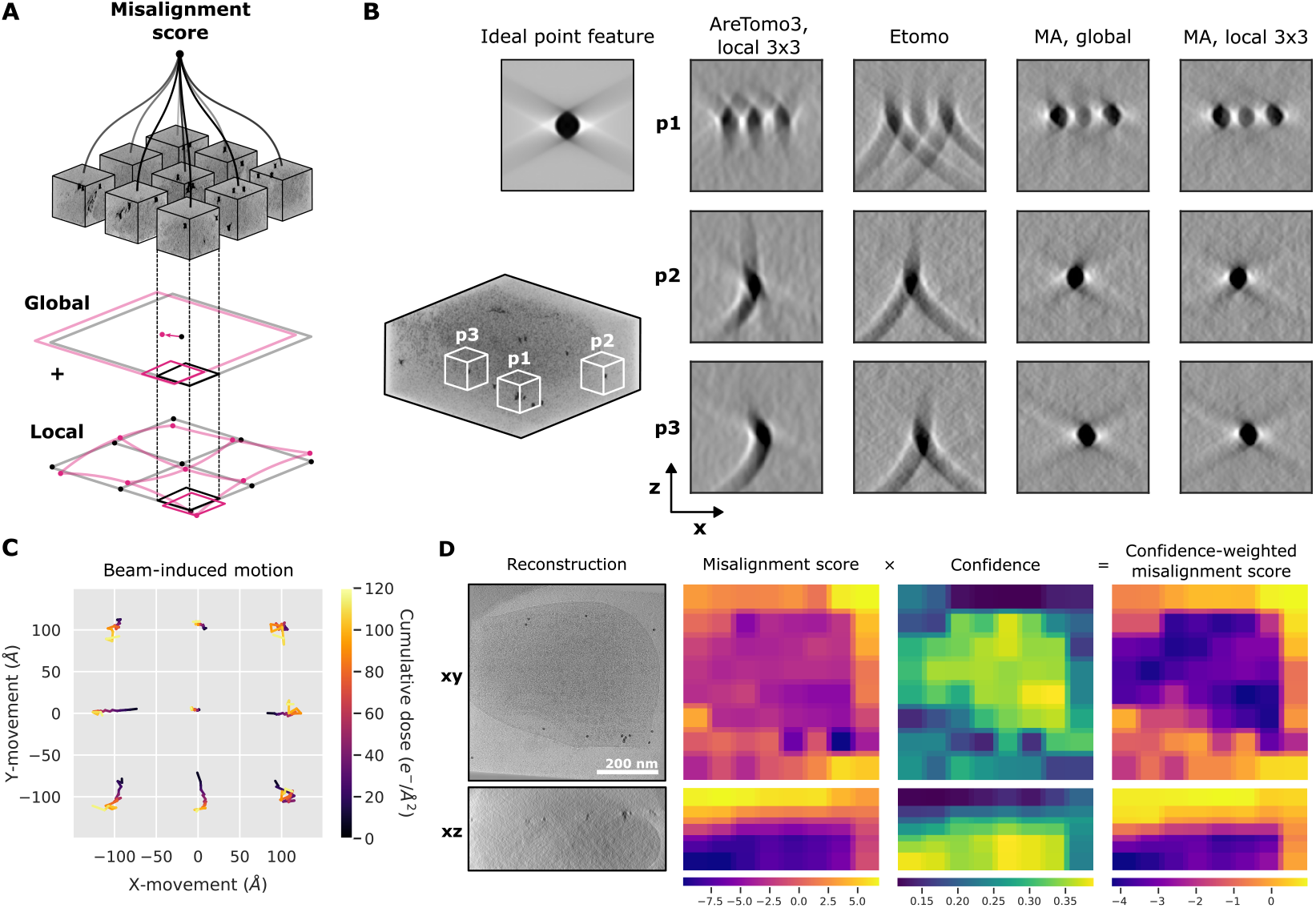
MissAlignment operates on small sub-volumes for local alignment while maintaining spatial attention over the field of view via a confidence mechanism. (A) Multiple sub-volumes are scored in batches over the full tomogram space. Each sub-volume can be corrected for global and local shifts via the projection model, illustrated here for a 3 ×3 image warp grid under orthographic projection along z. A deformation field with a temporal component can also be applied. (B) Reconstructions of gold-bead point features spread across one tomogram (EMPIAR-10499) illustrating local alignment quality across methods; subvolume p1 contains 3 gold beads; p2 and p3 each contain 1 gold bead. MA refers to MissAlignment. (C) Exposure-ordered 3×3 image warp grid parameters for positive tilt angles of a single tilt-series. Grid nodes are displaced by ±100 Å in y and x for visualization. Cumulative exposure is indicated from dark (first exposure) to light (last). (D) Merged predictions of misalignment score, confidence, and confidence-weighted misalignment score across all sub-volumes of one tomogram. xy- and xz-views are mean-intensity projections along z and y, respectively.

Specifically, MissAlignment uses a PyTorch ^20^ replica of the deformation model from Warp ^3^ and M ^6^, optimizing both global image shifts and “image warp” parameters. Because the implementation is fully differentiable, enabled by open-source libraries from teamtomo ^21^ and Warp and M ^22^, gradients are obtained efficiently. The resulting parameters are directly compatible with Warp and M for downstream analysis.

Scoring the full tomogram in a single pass is impractical given the memory demands of 3D CNNs. Instead, MissAlignment scores sub-volume reconstructions in small batches during both training and inference. At inference, image shift gradients are accumulated over all batches before each gradient-descent step (Fig. 2a). This sub-volume approach also enables computationally efficient Fourier-space reconstruction with local alignments, as deformation across each sub-volume is negligible.

The resulting alignment is accurate across the full tomogram, as demonstrated by the reconstruction of gold-bead point features (Fig. 2b). Ordered by accumulated exposure, the local shifts recapitulate beam-induced sample doming (Fig. 2c; ^1^).

### Confidence estimation as large-scale spatial attention

Empty sub-volumes can bias the overall misalignment score. Because sample thickness and orientation are not known a priori, some sub-volumes may contain no useful features for scoring. Conventional voxel-level spatial attention cannot solve this: MissAlignment scores independent sub-volumes, so entire volumes—not just voxels—may be unreliable, but are scored on the same scale as reliable sub-volumes. If the sample is thinner than assumed, scores from empty regions above and below the sample dominate and degrade accuracy.

MissAlignment addresses this with a second prediction head that estimates a confidence score for each sub-volume. During training, the model learns to down-weight the triplet loss for low-confidence examples, with a regularization term preventing collapse to zero confidence. This aspect of the training is also unsupervised.

At inference, misalignment scores are weighted by confidence before accumulation across the tomogram (Fig. 2d), effectively mimicking single-pass scoring with spatial attention. This makes MissAlignment robust to uneven sample distribution: in practice, high confidence maps to regions with biological features, while empty ice regions receive low confidence.

### MissAlignment rivals the accuracy of reference-based alignment

Reference-based alignment is the current gold standard for accuracy but requires large, homogeneous particles and a lengthy processing pipeline. Poor alignment propagates into all downstream steps, including template matching and 3D classification.

We benchmarked alignment accuracy by STA resolution after a standardized pipeline of template matching and pose optimization. MissAlignment in global-only (1×1) and local (3×3) mode was compared against automated reference-free methods and reference-based alignment. For 70S ribosomes (EMPIAR-10499 ^6^), particles were identified by template matching and pose-refined in M with frozen alignment parameters. Nucleosomes (EMPIAR-11830 ^23^) additionally required two rounds of 3D classification in RELION ^7^ to obtain clean particle sets.

Even in global-only (1 ×1) mode, MissAlignment improves resolution (10.4 Å nucleosomes, 9.3 Å ribosomes) over AreTomo3 (11.9 Å, 12.3 Å) and etomo patch-tracking (12.3 Å nucleosomes) or gold-fiducial alignment (10.6 Å ribosomes) (Fig. 3a–c). For ribosomes, the improvement partly reflects higher particle counts from more accurate template matching (Fig. 3d); for nucleosomes, particle counts are comparable and the resolution gain is driven by alignment accuracy alone. The absolute alignment error at high tilts (Fig. 1e) would predict a larger difference, but exposure weighting reduces the contribution of high-tilt images to the reconstruction.

**Figure 3:**
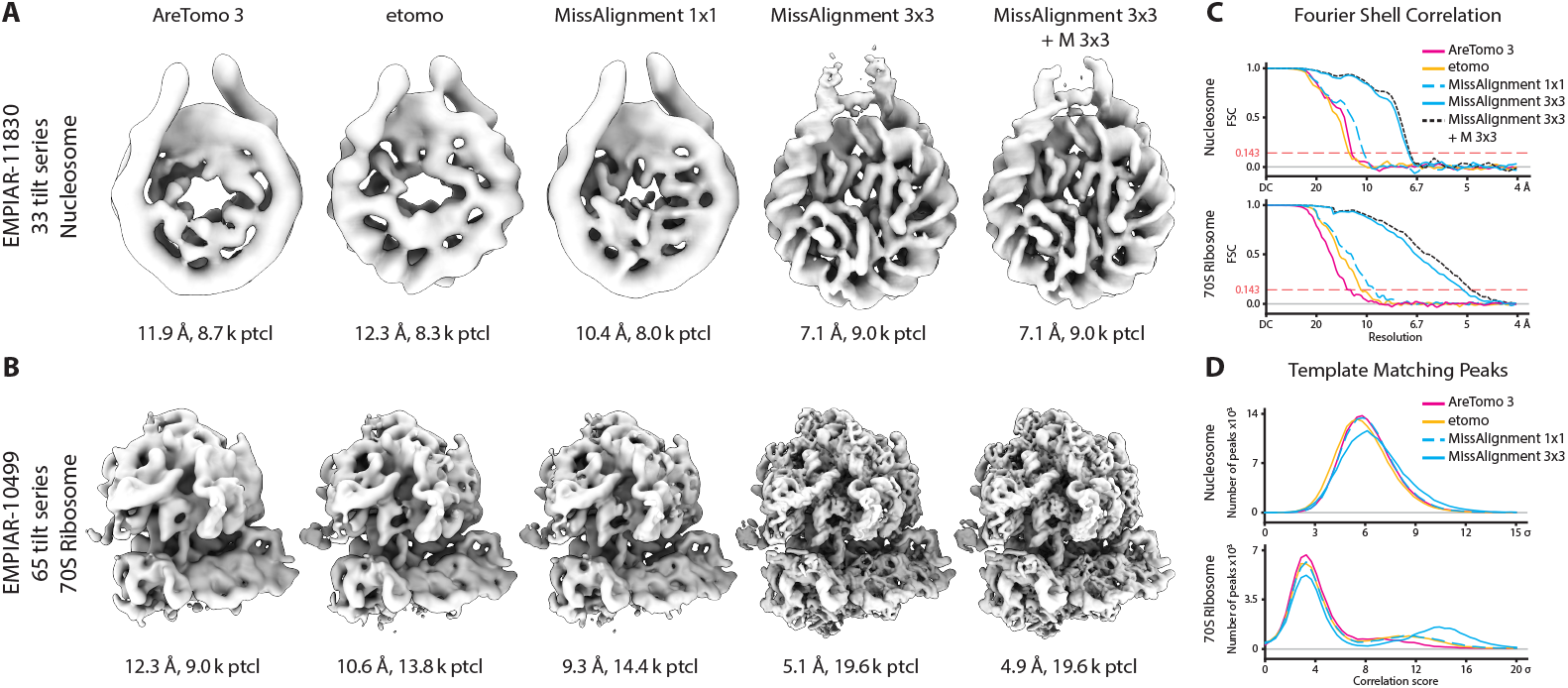
STA results from MissAlignment outperform fiducial-free methods and rival reference-based alignment. (A) Reconstructed maps with resolution and number of picked particles for nucleosomes from 33 tilt-series (EMPIAR-11830) after alignment with each method. Etomo uses patch tracking. (B) Same as (A) for 70S ribosomes from 65 tilt-series (EMPIAR-10499), except etomo uses gold-fiducial alignment. (C) Fourier shell correlation curves for maps in (A) and (B). (D) Distribution of template-matching correlation scores (in *σ* above background) for the top 5,000 (EMPIAR-11830) and 2,000 (EMPIAR-10499) peaks per tomogram after gradient-descent refinement.

**Figure 4:**
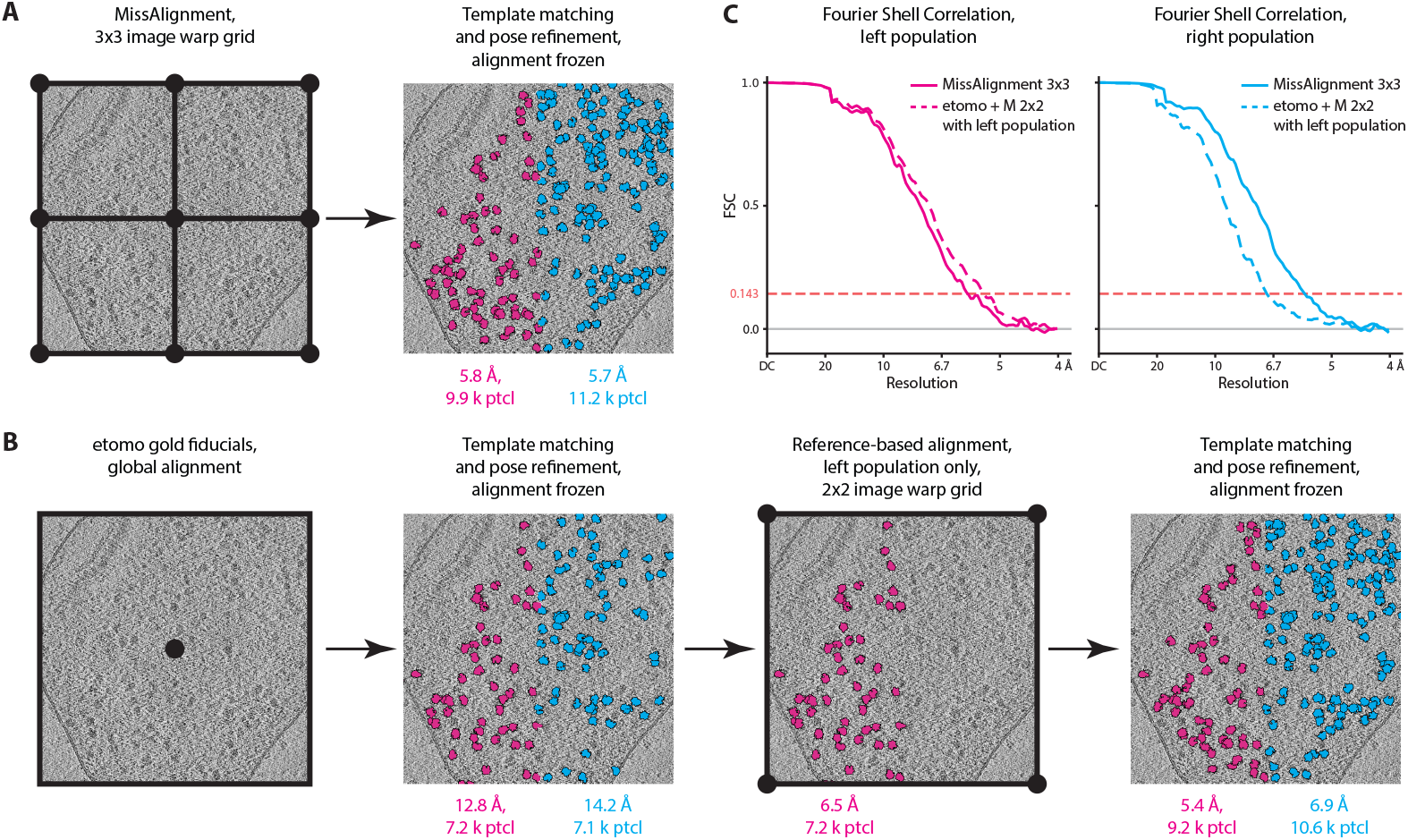
Local alignment in MissAlignment is not restricted to sample areas with reference structures. (A) STA results for 70S ribosomes in EMPIAR-10499 split into two sets based on the x-coordinate after 3 × 3 local alignment. (B) Data similarly split after gold-fiducial alignment in etomo. The left half was used for reference-based alignment in M with a 2 × 2 image warp grid; both halves were then repicked and pose-refined. (C) Fourier shell correlation curves for the left (pink) and right (blue) populations for MissAlignment 3 × 3 (solid) and etomo + M 2 × 2 (dashed).

**Figure 5:**
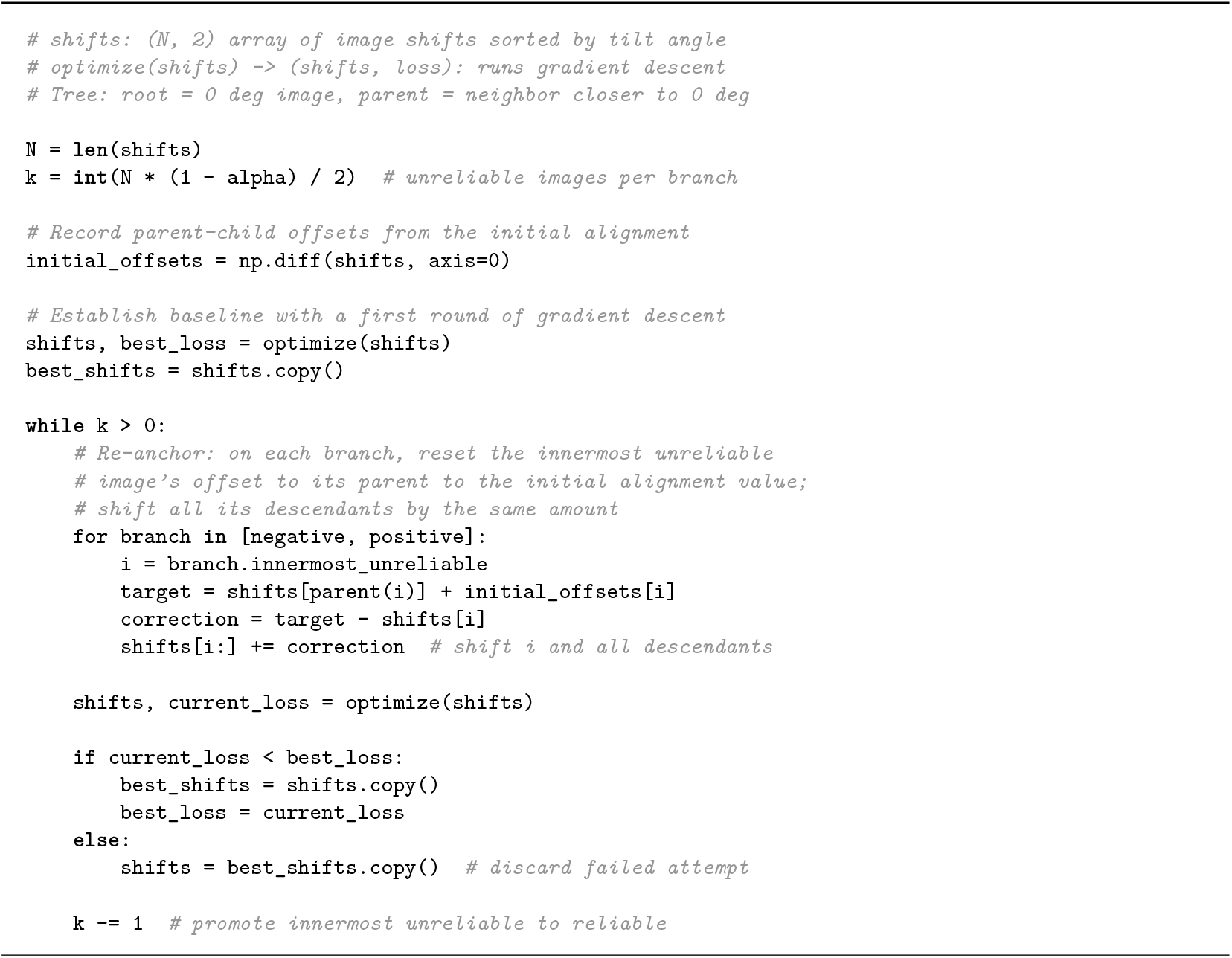
Algorithm 1. Iterative anchoring as Python-style pseudocode. The optimize function runs gradient descent on all tilt images and returns the optimized shifts together with the confidence-weighted misalignment score (eq. (4)).

Local alignment with a 3×3 image warp grid sub-stantially improves STA resolution over 1×1 alignment, reaching 7.1 Å for nucleosomes and 5.1 Å for ribosomes. Template-matching correlations also improve (Fig. 3d): for nucleosomes, overall scores increase; for ribosomes, a clear bimodal separation emerges between particles and background, and the number of detected particles rises from 14.4k to 19.6k (Fig. 3b).

To test whether reference-based refinement could improve upon these results, we used M to further optimize the 3 ×3 image warp parameters. For nu-cleosomes, resolution did not improve; for ribosomes, it improved marginally to 4.9 Å. MissAlignment thus rivals reference-based alignment accuracy without requiring a full processing pipeline.

### Full-tomogram alignment extends high-resolution analysis to new sample regions

When large particles such as ribosomes are available, reference-based alignment can use them to refine the initial alignment and improve analysis of nearby smaller targets. However, the two populations are not always co-localized—for example, when the target resides in an organelle devoid of ribosomes. The best available strategy is to fit a low-order deformation model (2×2 image warp) to the reference particles, since higher-order fits overfit in regions far from those particles, degrading accuracy below the reference-free baseline.

To compare this strategy against MissAlignment’s full-tomogram alignment, we divided the 70S ribosome population in EMPIAR-10499 into two spatially disjoint groups of equal size based on X coordinate. For reference-based alignment, only one group was used to optimize the image warp parameters, which were then frozen; the other group underwent pose refinement only. For MissAlignment, both groups underwent pose refinement only.

For the group used in reference-based alignment, resolution was slightly better (5.4 Å) than its MissAlignment counterpart (5.8 Å). However, the distant group fared worse (6.9 Å), whereas MissAlignment maintained comparable accuracy across both halves (5.7 Å). This disparity would grow with larger spatial separation between groups. Both strategies markedly outperformed gold fiducial-based alignment in etomo (12.8 Å and 14.2 Å for the same groups).

MissAlignment outperformed reference-based alignment in this common scenario while requiring no preliminary processing. By providing accurate alignment across the full field of view, MissAlignment opens new regions of existing datasets to high-resolution analysis and, for future data collection, removes the requirement for large reference particles—paving the path deep into large organelles such as the nucleus.

## Methods

### Software usage

MissAlignment is a reference-free tilt-series alignment method that integrates into cryo-ET processing pipelines at the alignment stage. It requires an initial alignment estimate from tools such as etomo ^12^ (patch-tracking or bead alignment) or AreTomo ^14^. Alternatively, a built-in cross-correlation step (torch-tiltxcorr) can generate the initial alignment, making MissAlignment fully standalone. This built-in initialization may yield slightly lower accuracy than etomo or AreTomo, particularly for thick specimens.

MissAlignment integrates most naturally into pipelines built around WarpTools ^3^, as it interfaces directly with the Warp/M metadata model. Its alignment parameters—including local deformation fields—can be used directly for tomogram reconstruction, template matching, and subtomogram averaging without conversion.

MissAlignment uses GPU acceleration for both training and alignment optimization, benefiting from multi-GPU systems where work is distributed across parallel workers, though single-GPU operation is also supported. The software processes an entire dataset at once through multiple coarse-to-fine iterations. Run-time scales with dataset size and iteration count; training time is relatively insensitive to dataset size, whereas alignment optimization scales more directly with the number of tilt series. For SHREC’21 (10 tilt series), total processing time on 4 NVIDIA RTX 4500 Ada GPUs was approximately 5.30 hours, of which ~5.17 hours was spent on training and ~13 minutes on alignment optimization.

### Configuring iterative refinement

The full MissAlignment pipeline alternates between CNN training and alignment optimization in a series of macro-iterations. In each iteration, the CNN is trained on the current alignment to learn a scoring function appropriate for the current misalignment level. The trained model is then used to optimize alignment parameters for all tilt series. The improved alignment serves as the starting point for the next iteration.

The iterations follow a coarse-to-fine progression: early iterations operate on downsampled data and optimize only global shifts, while later iterations use full-resolution data and optimize local deformation parameters. Because the CNN can extrapolate beyond its training distribution, each iteration improves the alignment further. However, the scoring function must be re-trained at progressively better alignment states, requiring multiple bootstrapping iterations to reach the optimum.

All hyperparameters are configurable via a .yaml file. The repository provides a default configuration used for multiple datasets in this study; exceptions are noted in the respective dataset sections below. MissAlignment supports “global”, “anchoring”, and “local” modes, detailed below.

### Contrastive training data generation

Because no ground-truth alignment exists, MissAlignment is trained contrastively using synthetic misalignment. For each training example, a sub-volume is reconstructed at a random position within a randomly selected tilt series using the current alignment parameters, producing an “aligned” example. A second reconstruction is calculated at the same position after adding a synthetic misalignment pattern to the current image shifts, producing a “misaligned” example. A third “anchor” sub-volume is a copy of either example, augmented with a different random mirror transformation to make it visually distinct. These three sub-volumes form a training triplet.

The synthetic misalignment is generated by composing four types of shift patterns, each applied independently with a configurable probability. *Trajectory* shifts produce smooth, parabolic drift across the tilt series, optionally with a breakpoint where the drift direction changes abruptly. *Jitter* adds independent Gaussian noise to each tilt image. *Outlier* shifts introduce a large sudden displacement on a single tilt. *Fracture* shifts apply a linear ramp or break to one end of the tilt series, simulating a strong error at one end of the tilt range. Multiple shift types can be combined in a single example, and at least one is always applied. All shifts are generated in 3D and then projected to 2D image shifts through the tilt geometry.

Each sub-volume is further augmented by mirroring along all axis combinations, random contrast scaling, masking of edge bands, and masking of random cubic regions within the volume.

### Misalignment scoring network

The default scoring network is a compact 3D CNN consisting of six convolutional layers, each followed by batch normalization and SiLU activation. The input is a single-channel sub-volume of 64^3^ voxels. Progressive stride-2 downsampling reduces the spatial dimensions through channel widths of 1, 8, 16, 32, 64, 64, and 64, followed by global average pooling to produce a 64-dimensional feature vector. A shared fully connected layer further reduces this to 16 features (with batch normalization and SiLU activation).

Two independent linear heads operate on the shared features. The score head (without bias) produces a scalar misalignment score, where lower values indicate better alignment. The log-precision head produces a scalar log-precision estimate, representing the model’s confidence in its score for a given sub-volume. The role of the precision head is described below.

### Precision-weighted triplet loss

MissAlignment is trained with a modified triplet margin ranking loss that incorporates the model’s precision estimates. Each triplet consists of two sub-volumes with the same alignment quality (the “close” pair: both aligned, or both misaligned, chosen randomly with equal probability) and one with different alignment quality (the “distant” member). Let *s*_*i*_ and *π*_*i*_ = exp(log *π*_*i*_) denote the predicted score and precision for the *i*-th triplet member. The positive distance *d*^+^ = |*s*_1_ −*s*_2_|measures the score discrepancy within the close pair, while the negative distance *d*^*−*^ measures the separation from the distant member. The per-triplet loss is:

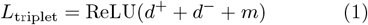

where *m* is the margin (by default, 0.5). Each triplet’s loss is weighted by the geometric mean of the three precisions, *w* = (*π*_1_· *π*_2_ ·*π*_3_)^1*/*3^, so that triplets involving sub-volumes in uninformative regions (e.g. empty space outside the sample) contribute less to the gradient. To prevent the precision estimates from collapsing to zero, a regularization term penalizes low precision:

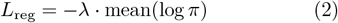

The total loss is the mean of the precision-weighted triplet losses plus the regularization term:

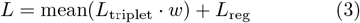

### Training procedure

The CNN is trained using AdamW ^24^ with a learning rate of 10^*−*3^ and weight decay of 10^*−*4^. Training proceeds for a maximum of 30 epochs per macro-iteration, where each epoch consists of 1000 steps with a batch size of 32 and sub-volume size of 96^3^. A linear warmup over the first 500 steps is followed by a multi-step decay that halves the learning rate at epochs 5 and 15. Early stopping monitors the training loss but activates only after the final learning rate reduction, ensuring the model benefits from the full schedule before convergence is assessed. The best checkpoint by training loss is used for the subsequent alignment step.

### Alignment optimization

After training, the model weights are frozen and the 3D CNN serves as the objective function for alignment optimization, which uses L-BFGS with Strong Wolfe line search.

3D CNNs are memory-intensive, making it impractical to score the full tomogram in a single forward pass. Instead, MissAlignment operates on cubic sub-volumes. A regular grid of overlapping positions is generated to cover the tomogram, and at each position, a sub-volume is reconstructed from the tilt images using the current alignment parameters.

The reconstruction is performed in Fourier space with 2×oversampling, following the same procedure used in Warp and M ^3;6^. The entire reconstruction pipeline is implemented in PyTorch ^20^ with differentiable operators, so gradients flow from the misalignment score through the reconstruction to the alignment parameters via automatic differentiation. Warpylib, a PyTorch replica of Warp’s algorithms, handles tiltseries I/O and differentiable reconstruction. It uses torch-subpixel-crop ^21^ for patch extraction and torch-projectors ^22^ for compiled differentiable forward and backward projection kernels.

Each reconstructed sub-volume is normalized to zero mean and unit variance before scoring. Because sample deformation is negligible over the spatial extent of a single sub-volume, the local rigid-body approximation holds, and the Fourier-space reconstruction remains accurate even when the global alignment model includes non-rigid deformation terms.

Because the tomogram is scored as a set of independent sub-volumes, gradients for the alignment parameters are accumulated across all sub-volume batches spanning the tomogram before each optimizer step. Each sub-volume’s contribution to the total loss is weighted by its predicted precision, so that the optimization is driven primarily by informative regions of the tomogram:

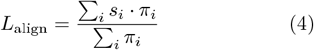

This confidence-weighted accumulation mimics single-pass scoring with spatial attention, making the optimization robust to variable sample thickness and distribution.

### Alignment parameterization

MissAlignment uses a PyTorch re-implementation of the deformation model employed by Warp and M ^3;6^, supporting three levels of increasing complexity. At the coarsest level, the model optimizes a single 2D shift per tilt image, correcting global tracking errors. At the next level, a cubic B-spline grid is defined over the spatial extent of each tilt image, with the grid values representing local 2D shifts. This “image warp” captures the spatially resolved component of beam-induced sample motion. At the finest level, a cubic B-spline grid is defined over the full 3D volume and over time (tilt index), representing a volumetric deformation field. Importantly, image warp is applied after projecting 3D positions into the 2D images, while volume deformation is applied before projections. Thus, volume warping directly models physical sample deformation.

Because the parameterization is identical to that of Warp and M, optimized parameters can be used directly in these programs without loss of accuracy.

After each optimization step, the alignment is recentered by projecting the global x- and y-shift at the zero-tilt image back through the tilt geometry to all other images. This ensures that the x- and y-origin of the reconstruction remain consistent and prevents the optimization from introducing a global translational drift.

### Alignment with iterative anchoring

Correlation-based alignment provides accurate relative shifts between adjacent images, but these accumulate into large errors across the full tilt series because virtually no information overlap constrains non-adjacent tilts. When gradient descent operates on such an alignment, it must simultaneously correct small relative errors between adjacent tilts and large accumulated errors at the extremes of the tilt range. This can cause the optimizer to reduce errors within clusters of adjacent images at the cost of introducing abrupt alignment breaks between clusters. We address this with a procedure called *iterative anchoring*.

Iterative anchoring treats the angle-sorted tilt series as a tree with two branches rooted at the central tilt image: one branch extends toward the most negative tilt angle, the other toward the most positive (see Algorithm 1). In this tree, each image’s *parent* is its neighbor closer to 0° and its *child* is its neighbor further from 0°. The central images (by default, the inner half of the tilt range) are designated “reliable”, meaning gradient descent is unlikely to introduce breaks between them. The remaining images on each branch are designated “unreliable”.

Gradient descent is first run without any anchoring to establish a baseline solution. Then, at the start of each subsequent iteration, the innermost unreliable image on each branch is re-anchored: its relative offset to its reliable parent is reset to the value from the initial correlation-based alignment. All unreliable descendants keep their relative shifts, i.e. their absolute shifts are adjusted by the same amount. Gradient descent is then run on the full tilt series. If the resulting loss improves upon the current best, the new alignment is accepted; otherwise the previous best is restored. Regardless of the outcome, the innermost unreliable image on each branch is then promoted to reliable, and the process repeats until all images are marked reliable.

In practice, iterative anchoring is applied during the early macro-iterations of MissAlignment (see above), when coarser downsampling factors (e.g., 3× or 2×) make repeated gradient descent over the full reconstruction computationally cheap.

### Dataset preprocessing and evaluation SHREC-2021

All tilt-series from the SHREC-2021 dataset ^19;25^ were preprocessed by downsampling the images to 10 Å via cropping in Fourier space. Images, ground-truth alignments, and torch-tiltxcorr alignments were stored as Python pickle files in a Zenodo archive ^26^. A repository script (`examples/shrec/preproc.py`) downloads and converts these files to a Warp-compatible format.

MissAlignment was benchmarked against Are-Tomo2^14^ on the SHREC-2021 dataset. Both programs were run without local alignment, as the simulation did not include beam-induced motion. AreTomo2 was executed on the 10 Å tiltstack, keeping the tilt-axis fixed (by specifying `-TiltAxis 0.0001 -1,` to circumvent a known issue where specifying 0.0 causes unwanted optimization), turning off pretilt estimation, and fixing the alignment z-height (`-AlignZ`) to 180 voxels. The dataset was simulated with identical sample box dimensions and no tilt-axis or tilt-angle offset, so these fixed values give AreTomo2 optimal performance.

MissAlignment was run in standalone mode with initial alignment provided by torch-tiltxcorr without pretilt estimation. The program was configured to run 6 macro-iterations, with the first two operating on 2×downsampled data (20 Å) with iterative anchoring, and the remaining four at 10 Å with global-only alignment. For comparison, the final alignments from both programs were placed into the ground-truth reference frame. Because alignments can accumulate drift along z, tomograms were reconstructed with Warpylib and the 3D shift to the ground truth was estimated by cross-correlation. This shift was projected to per-tilt 2D offsets via the tilt geometry and subtracted from the estimated alignments. Misalignment was quantified as the absolute distance to the ground truth per tilt and averaged across tilts per tilt-series.

### EMPIAR-10499

The dataset was preprocessed in WarpTools with settings similar to those described previously ^6^.

Tilt-series alignment with gold beads in IMOD/etomo ^12^ and with AreTomo3^15^ was performed automatically via WarpTools wrappers at a pixel size of 10 Å. For etomo, the tilt-axis angle was first optimized across all tilt-series, and the median was selected as a fixed parameter for a subsequent run. AreTomo3 was run with the same fixed tilt-axis angle. Sample thickness was detected automatically. AreTomo3 was run with 3 × 3 local patches for visual inspection of local alignment on gold beads. For subtomogram averaging, AreTomo3 alignment was performed in global-only mode, as its local alignments are not, to our knowledge, compatible with any subtomogram averaging workflow.

Alignment with MissAlignment was optimized over 8 macro-iterations: 2 coarse iterations at 20 Å in “iterative anchoring” mode, 1 fine iteration at 10 Å with global-only alignment, and 5 iterations of local alignment using a 3 ×3 image warping grid. The global-only alignment results after iteration 3 were used to evaluate the global alignment performance of MissAlignment, while the local alignments from the final iteration were used for subtomogram averaging with local alignments. For further processing, template matching in Warp-Tools was performed across all alignment conditions at a pixel size of 10 Å using EMD-11650 as template. A uniform threshold was applied across all conditions, yielding varying particle counts depending on alignment quality. Particle poses were first refined using a template-matching algorithm in WarpTools that performs a local pose search for annotated peaks via gradient descent, then further refined in M without fitting any tilt-series alignment parameters. The resulting half-maps were used for gold-standard FSC calculation to determine resolution, and the denoised maps were used for visualization.

The 3×3 local alignment results from MissAlignment were then subjected to an additional round of M refinement with optimization of the 3×3 image warping grid parameters enabled, providing a direct comparison to alignments obtained by reference-based methods.

For the population-split experiment, particles were initially picked by template matching on the etomo-aligned tomograms. Particles were divided into two sets based on their x-coordinate, with the split at the tomogram center. The alignment was first refined with MissAlignment using a 3×3 local image warping grid, after which the poses of both particle sets were refined with M and used to determine resolution. Separately, the etomo alignment was refined in M with a 2 ×2 image warping grid using only particles from the positive half. With this alignment frozen, pose refinement was then performed on the negative half, and the resulting reconstruction was used to determine resolution.

### EMPIAR-11830

Raw EER exposures, MDOC files, and gain references for 33 tilt-series containing nucleosomes ^23^were downloaded from EMPIAR (Table S1). The dataset was preprocessed in WarpTools.

Alignment scenarios analogous to EMPIAR-10499 were explored: etomo patch-tracking, AreTomo3 in global-only mode, MissAlignment with 1 ×1 and 3×3 grids, and M refinement with a 3 ×3 grid following the local alignments from MissAlignment. Template matching was unreliable for nucleosomes due to their small size, requiring an alternate strategy. Particles were liberally picked and subjected to multiple rounds of 3D classification. The first round used 6 distinct classes obtained by running initial model generation in RELION using MissAlignment’s alignments. The second round used particles from the best class of the first round, and 6 copies of the corresponding map.

### Sample preparation, cryo-FIB milling, and cryo-ET of ovarian cortex cells

MissAlignment was also tested on tilt series from bovine ovarian cells (Fig. S3). Fresh cow ovaries were provided by the Utrecht University Veterinary Faculty. The ovarian cortex (the outermost ∼1-mm layer) was dissected and cut into ∼2-mm pieces. Samples were digested with Liberase™ Research Grade (Roche) at a final concentration of 0.39 Wunsch Units/mL in L-15 medium (Sigma) at 37°C for a total of 75 minutes with gentle trituration every ∼15 minutes. Digestion was terminated by adding 10% fetal bovine serum (FBS) in a 1:1 volume ratio. Fragmented samples were successively passed through 300-µm, 100-µm, 60-µm, and 30-µm Nylon mesh filters (pluriSelect). Material collected on the 30-µm mesh was retrieved by inverting the filter and flushing with 5 mL of DPBS (Sigma). Isolated ovarian cortex cells were successively incubated in a cryoprotectant gradient of ethylene glycol (EG) and dimethyl sulfoxide (DMSO) at concentrations of 1.75%/1.75%, 2.33%/2.33%, 3.5%/3.5%, and finally 7.5%/7.5%. Approximately 4 µL of cell suspension was pipetted onto glow-discharged Quantifoil R2/1 200-mesh holey carbon grids and blotted from the back for ∼6 s with Whatman #1 filter paper using a manual plunger (MPI Martinsried). Grids were then immediately plunged into an ethane (37%)/propane mix cooled to liquid nitrogen temperatures.

Grids were clipped into autogrid frames (ThermoFisher) and loaded into the Aquilos 2 (ThermoFisher) at the Utrecht University Electron Microscopy Centre (UU-EMC). Grids were sputter coated (30 mA current for 10 s) and a protective GIS layer deposited for ∼15 s. Lamella sites were milled to a thickness of ∼2 µm using a 0.5 nA beam current, then to ~600 nm with a 0.3 nA current, and to ∼250–300 nm using a 0.1 nA current. These steps were automated with AutoTEM (ThermoFisher). Polishing was performed manually to a final target thickness of ∼200 nm using a 30 pA current.

Cryo-ET data were acquired on the UU-EMC Talos Arctica (ThermoFisher) operating at 200 kV and equipped with a K2 direct electron detector and energy filter (Gatan). Tilt series were collected at a nominal magnification of 49 kX (unbinned pixel size of 2.83 Å) with a grouped dose-symmetric tilt scheme 27 covering a range from −35° to +45° in 3° increments. The total exposure was ∼ 118 e*−*/Å ^2^.

WarpTools was used for dataset preprocessing. Motion correction was performed with 1×1×13 parameters for each tilt’s frame series, and a 2× 2 parameter grid for CTF estimation. For each frame, even and odd exposure halves were written out for tomogram denoising. Tilt-series alignment was performed on 10 Å stacks using both AreTomo3 and MissAlignment. AreTomo3 was run with CTF estimation disabled, a starting volume height of 500 pixels, and without local patch alignment, as the latter worsened results upon visual inspection. MissAlignment was run with settings similar to those used for EMPIAR-10499, except that the first coarse iteration was performed at 30 Å instead of 20 Å.

The MissAlignment results were used directly in WarpTools to reconstruct even and odd CTF-corrected tomograms. AreTomo3 results were first imported into Warp’s metadata format before reconstructing half-maps. Tomograms were denoised with Noise2Map in WarpTools using default parameters and 15,000 training iterations. The denoised tomograms were used for visualization. Full tomograms were mutually aligned by 3D cross-correlation and interpolated using *n*-dimensional spline interpolation in SciPy28. Cropped regions were additionally aligned locally to account for shifts introduced by MissAlignment’s local alignment.

### Data visualization

All plots and graphs were generated with matplotlib 29 and seaborn 30. Views of tilt images and 2D slices and projections of tomograms were rendered using spline interpolation in matplotlib. 3D views of tomograms were generated in Napari 31. Loading tomography data into Python was handled with mrcfile 32, while typical processing and filtering operations were handled with NumPy 33 and SciPy 28. Most image and volume shifting operations were performed with torch-fourier-shift 21.

## Discussion

MissAlignment is a machine-learning approach to tilt-series alignment that requires no ground-truth training data. Starting from an imperfect alignment, it iteratively bootstraps toward an improved solution in a fully unsupervised manner, while a confidence weighting mechanism focuses the model on the most informative sample regions. To our knowledge, this is the first method to replace fixed similarity metrics with a learned scoring function for de novo tilt-series alignment. MissAlignment achieves local alignment accuracy rivaling reference-based methods early in the processing pipeline, saving considerable time, and extends accurate subtomogram averaging to samples that lack the large reference structures these methods require.

STA resolution benefits most from local alignment, while global-only alignment contributes modestly. Yet global-only alignment produces strikingly sharper reconstructions of point features such as gold beads. Because low-angle tilts are acquired early in the exposure, they retain the most high-resolution signal and dominate STA resolution. Existing tools already align these images reasonably well; errors accumulate mainly at high tilts (Fig. 1e), where they distort reconstructions but contribute little to high-resolution STA. Consistent alignment across all tilts nonetheless benefits analyses that depend on low-resolution signal throughout the tilt range, such as segmentation. Indeed, for *in situ* lamellae, MissAlignment reconstructions show improved interpretability of low-abundance structures and rare events (Fig. S3).

MissAlignment can be applied to any dataset without manual intervention, but it does not make reference-based refinement obsolete. M can additionally refine CTF parameters and stage angles to reach 3.5 Å on the same ribosome particles. MissAlignment provides an accurate starting point for such refinement, and enables high-resolution STA on samples where reference-based methods are not applicable.

Accuracy and speed can be further improved by guiding reconstruction location selection. The confidence head already focuses the model on informative regions, but automatic sample boundary estimation or segmentation could complement it by restricting training and alignment to biologically relevant areas— particularly when ice contamination or preparation artifacts are present.

The core principle underlying MissAlignment— learning to distinguish well-aligned from degraded reconstructions via synthetic perturbations, then optimizing parameters against the learned score—extends naturally beyond tilt-series alignment. Within cryo-EM and cryo-ET, any parameter estimation step that relies on a fixed similarity metric could benefit from a learned objective, including beam-induced motion correction, defocus estimation, or projection geometry refinement.

An exciting prospect is whether trained models can generalize across datasets, eliminating the need for perdataset training. Model training currently accounts for most of MissAlignment’s runtime, yet the learned features may well transfer. Compiling a diverse training set from public repositories such as EMPIAR-11830 23 could yield a general-purpose scoring function that works out of the box, making high-quality alignment accessible as a routine step in any cryo-ET pipeline.

## Data and Code Availability

MissAlignment is available under a BSD-3 License:

https://github.com/warpem/miss-alignment.

## Acknowledgements

We thank Przemysław Dutka from Genentech, Maurice Frijns and Sander Roet from Utrecht University, and Josh Hutchings and Utz Ermel from CZI Biohub for beta-testing MissAlignment. The authors used Claude (Anthropic) to assist with writing, editing, and coding. All AI-generated content was reviewed and verified by the authors, who take full responsibility for the final content.

## Contributions

A.B., M.L.C., and D.T. designed and implemented the MissAlignment software. M.L.C. and D.T. bench-marked the software against the SHREC’21 dataset.

M.L.C. and D.T. tested and evaluated alignment performance on ribosomes. D.T. evaluated alignment performance on nucleosomes. J.v.L. and M.R.L. prepared and collected tilt series of ovarian cortex cells lamellae, and M.L.C. and J.v.L. processed the data.

D.T. provided the idea and project design. D.T. and M.R.L. provided funding. M.R.L. provided resources and additional interpretation. A.B., M.L.C., M.R.L., and D.T. wrote the manuscript.

## Competing Interests

The authors declare no competing interests.

---

**Figure S1:**
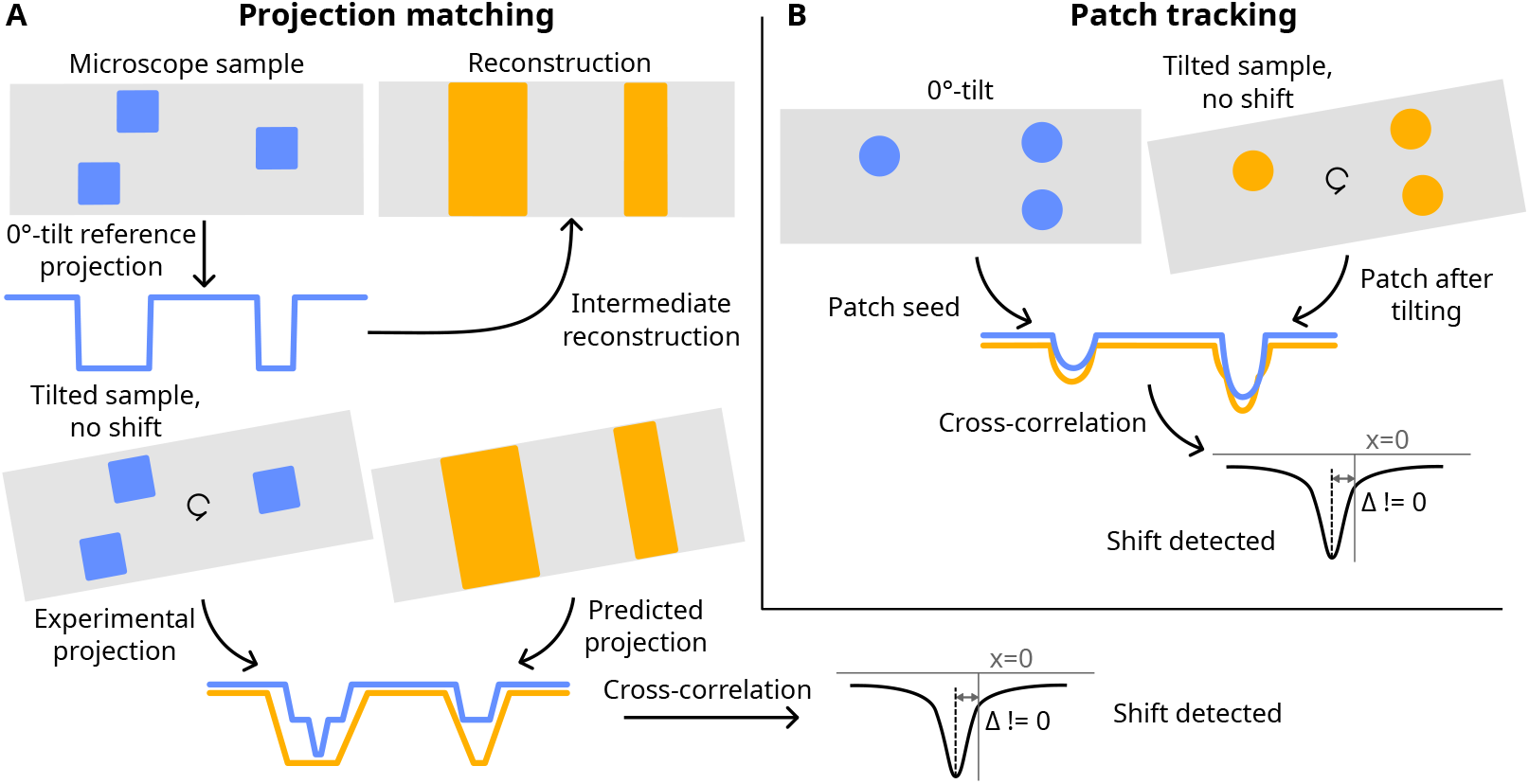
Failure modes of fiducial-free alignment methods, illustrated by projecting a 2D sample to 1D line profiles. (A) Projection matching fails when density is unevenly distributed. Progressive alignment from 0° to higher tilts causes mismatch between experimental and predicted projections, leading to error accumulation.(B) Patch-tracking cross-correlation incorrectly detects an image shift when multiple objects overlap along z and are unevenly distributed.

**Figure S2:**
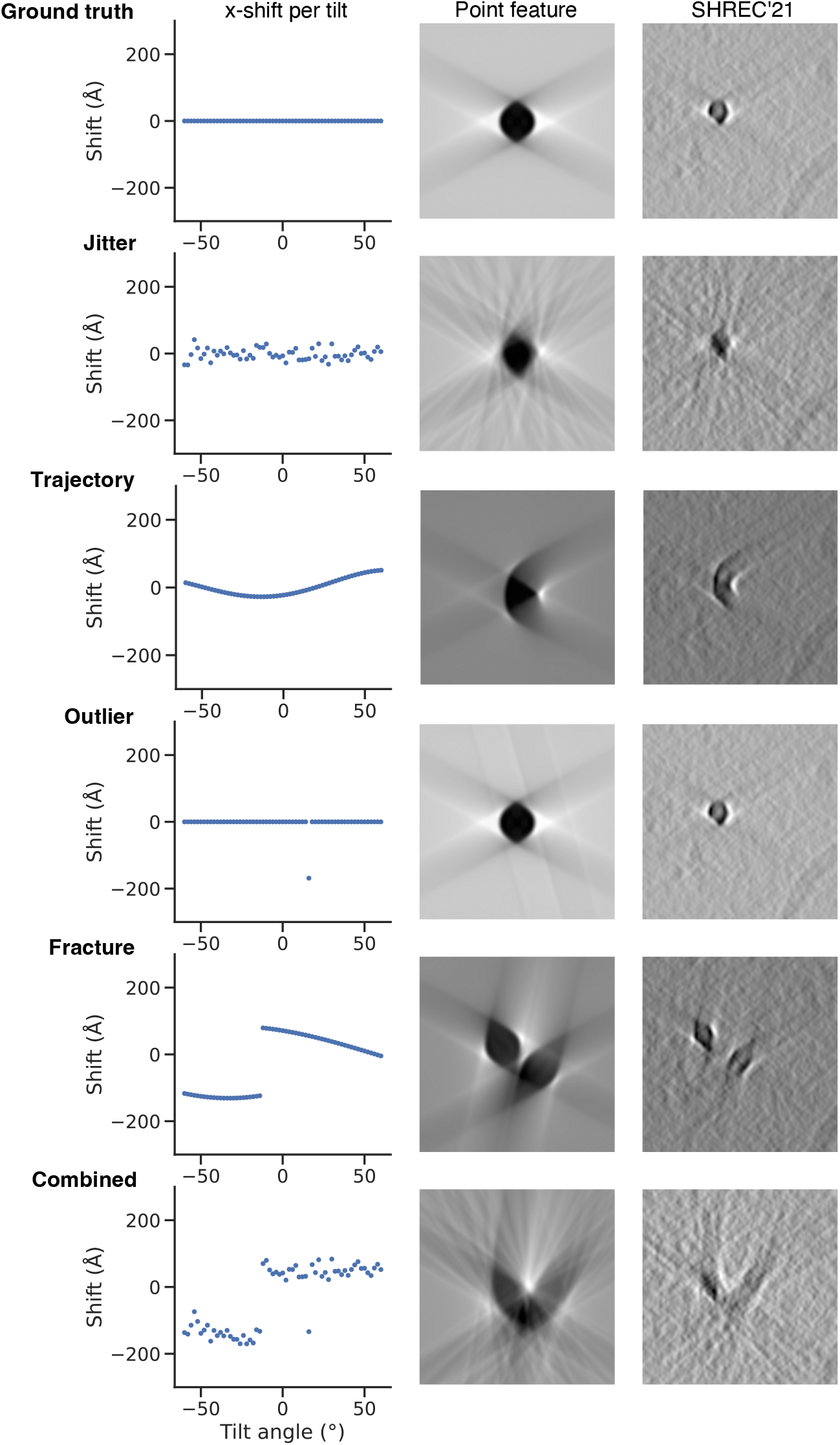
Synthetic misalignment types used for generating contrastive training examples. Each row shows a different perturbation type. Left: x-shift (perpendicular to the tilt axis) per tilt angle. Middle: effect on an ideal point-feature reconstruction. Right: effect on a reconstructed SHREC’21 patch.

**Figure S3:**
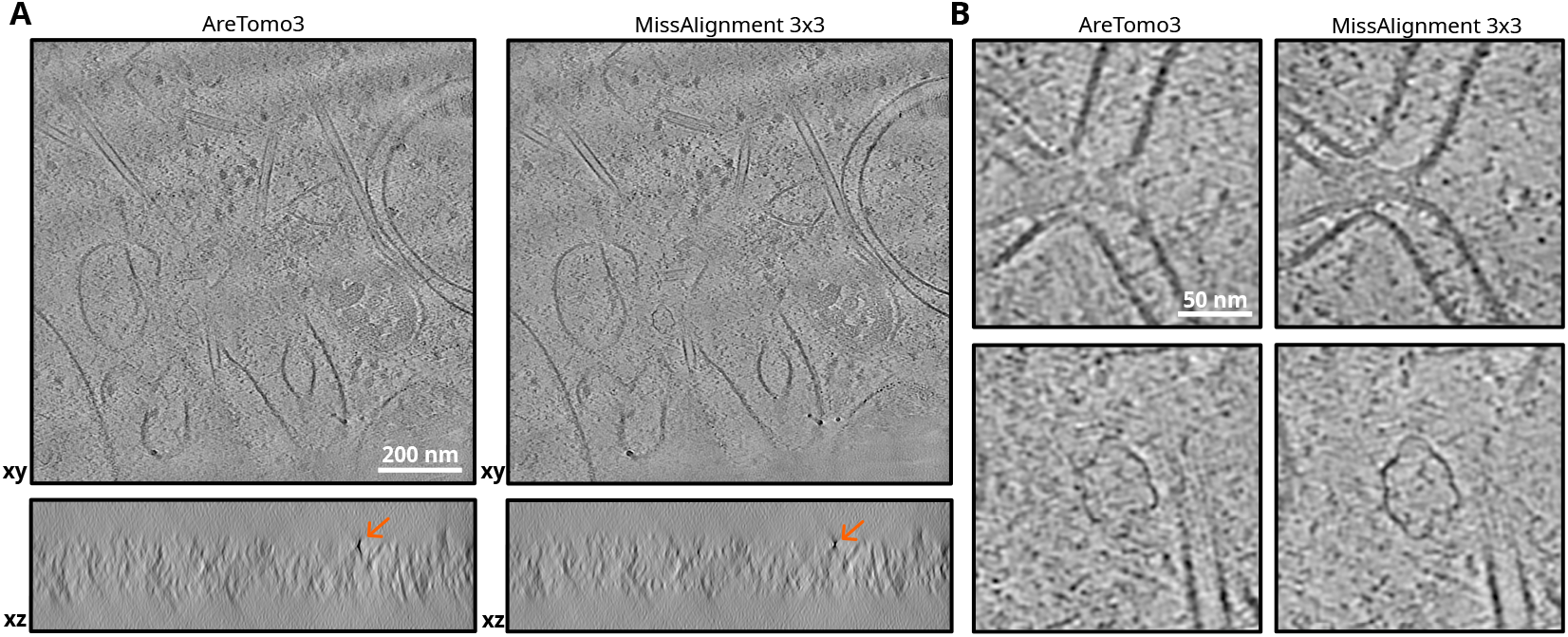
MissAlignment improves visualization of rare biological features *in situ* compared to AreTomo3. (A) Full-view xy- and xz-slices of an ovarian cortex cell lamella. Orange arrow on the xz-slice indicates a point feature highlighting the difference in alignment quality. (B) Zoomed-in views of two low-abundance structures where MissAlignment improves contrast.

**Table S1:**
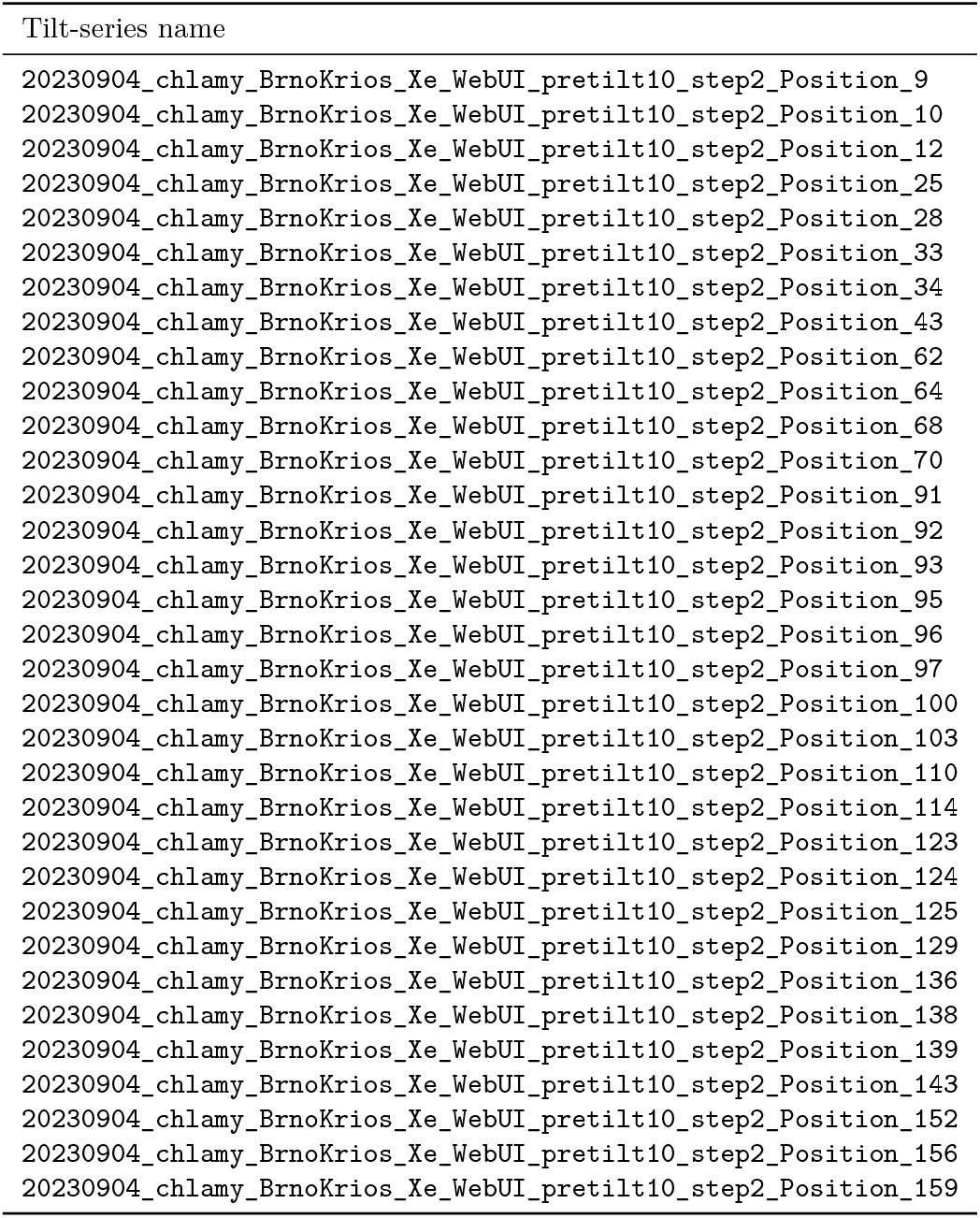
Tilt-series from EMPIAR-11830 used in this study.

| Tilt-series name |
| --- |
| 20230904_chlamy_BrnoKrios_Xe_WebUI_pretilt10_step2_Position_9 |
| 20230904_chlamy_BrnoKrios_Xe_WebUI_pretilt10_step2_Position_10 |
| 20230904_chlamy_BrnoKrios_Xe_WebUI_pretilt10_step2_Position_12 |
| 20230904_chlamy_BrnoKrios_Xe_WebUI_pretilt10_step2_Position_25 |
| 20230904_chlamy_BrnoKrios_Xe_WebUI_pretilt10_step2_Position_28 |
| 20230904_chlamy_BrnoKrios_Xe_WebUI_pretilt10_step2_Position_33 |
| 20230904_chlamy_BrnoKrios_Xe_WebUI_pretilt10_step2_Position_34 |
| 20230904_chlamy_BrnoKrios_Xe_WebUI_pretilt10_step2_Position_43 |
| 20230904_chlamy_BrnoKrios_Xe_WebUI_pretilt10_step2_Position_62 |
| 20230904_chlamy_BrnoKrios_Xe_WebUI_pretilt10_step2_Position_64 |
| 20230904_chlamy_BrnoKrios_Xe_WebUI_pretilt10_step2_Position_68 |
| 20230904_chlamy_BrnoKrios_Xe_WebUI_pretilt10_step2_Position_70 |
| 20230904_chlamy_BrnoKrios_Xe_WebUI_pretilt10_step2_Position_91 |
| 20230904_chlamy_BrnoKrios_Xe_WebUI_pretilt10_step2_Position_92 |
| 20230904_chlamy_BrnoKrios_Xe_WebUI_pretilt10_step2_Position_93 |
| 20230904_chlamy_BrnoKrios_Xe_WebUI_pretilt10_step2_Position_95 |
| 20230904_chlamy_BrnoKrios_Xe_WebUI_pretilt10_step2_Position_96 |
| 20230904_chlamy_BrnoKrios_Xe_WebUI_pretilt10_step2_Position_97 |
| 20230904_chlamy_BrnoKrios_Xe_WebUI_pretilt10_step2_Position_100 |
| 20230904_chlamy_BrnoKrios_Xe_WebUI_pretilt10_step2_Position_103 |
| 20230904_chlamy_BrnoKrios_Xe_WebUI_pretilt10_step2_Position_110 |
| 20230904_chlamy_BrnoKrios_Xe_WebUI_pretilt10_step2_Position_114 |
| 20230904_chlamy_BrnoKrios_Xe_WebUI_pretilt10_step2_Position_123 |
| 20230904_chlamy_BrnoKrios_Xe_WebUI_pretilt10_step2_Position_124 |
| 20230904_chlamy_BrnoKrios_Xe_WebUI_pretilt10_step2_Position_125 |
| 20230904_chlamy_BrnoKrios_Xe_WebUI_pretilt10_step2_Position_129 |
| 20230904_chlamy_BrnoKrios_Xe_WebUI_pretilt10_step2_Position_136 |
| 20230904_chlamy_BrnoKrios_Xe_WebUI_pretilt10_step2_Position_138 |
| 20230904_chlamy_BrnoKrios_Xe_WebUI_pretilt10_step2_Position_139 |
| 20230904_chlamy_BrnoKrios_Xe_WebUI_pretilt10_step2_Position_143 |
| 20230904_chlamy_BrnoKrios_Xe_WebUI_pretilt10_step2_Position_152 |
| 20230904_chlamy_BrnoKrios_Xe_WebUI_pretilt10_step2_Position_156 |
| 20230904_chlamy_BrnoKrios_Xe_WebUI_pretilt10_step2_Position_159 |

## Notes

https://github.com/warpem/miss-alignment

### Competing Interest Statement

The authors have declared no competing interest.

